# μ-Theraphotoxin-Pn3a inhibition of Ca_V_3.3 channels reveals a novel isoform-selective drug binding site

**DOI:** 10.1101/2021.10.04.463006

**Authors:** Jeffrey R. McArthur, Jierong Wen, Andrew Hung, Rocio K. Finol-Urdaneta, David J. Adams

## Abstract

Low voltage-activated calcium currents are mediated by T-type calcium channels Ca_V_3.1, Ca_V_3.2, and Ca_V_3.3, which modulate a variety of physiological processes including sleep, cardiac pace-making, pain, and epilepsy. Ca_V_3 isoforms’ biophysical properties, overlapping expression and lack of subtype-selective pharmacology hinder the determination of their specific physiological roles in health and disease. Notably, Ca_V_3.3’s contribution to normal and pathophysiological function has remained largely unexplored. We have identified Pn3a as the first subtype-selective spider venom peptide inhibitor of Ca_V_3.3, with >100-fold lower potency against the other T-type isoforms. Pn3a modifies Ca_V_3.3 gating through a depolarizing shift in the voltage dependence of activation thus decreasing Ca_V_3.3-mediated currents in the normal range of activation potentials. Paddle chimeras of K_**V**_1.7 channels bearing voltage sensor sequences from all four Ca_V_3.3 domains revealed preferential binding of Pn3a to the S3-S4 region of domain II (Ca_V_3.3^DII^). This novel T-type channel pharmacological site was explored through computational docking simulations of Pn3a into all T-type channel isoforms highlighting it as subtype-specific pharmacophore with therapeutic potential. This research expands our understanding of T-type calcium channel pharmacology and supports the suitability of Pn3a as a molecular tool in the study of the physiological roles of Ca_V_3.3 channels.

## Introduction

Voltage-gated calcium (Ca_V_) channels are activated by membrane depolarization and are involved in a number of physiological processes including contraction, secretion, neurotransmitter release, and gene expression (Catterall et al. 2005). In contrast to high voltage-activated (HVA) Ca^2+^ currents mediated by L-, N-, P/Q- and R-type calcium channels that require large depolarisations, low voltage-activated (LVA) currents mediated by T-type Ca^2+^ channels are activated by small membrane depolarization and display distinctively faster activation and inactivation kinetics. T-type channel activation near resting membrane potentials generate low-threshold Ca^2+^ spikes responsible for the conspicuous burst firing and low-frequency oscillatory discharges observed in thalamic, olivary, and cerebellar neurons (Park et al. 2010, Dreyfus et al. 2010, Molineux et al. 2006, Lee et al. 2014). Thus, the biophysical properties of the T-type channels make them important regulators of cardiac and neuronal excitability (Perez-Reyes 2003, Huguenard 1996, Coulter, Huguenard, and Prince 1989) and therefore key pharmacological targets for the treatment of neurological and psychiatric disorders (Zamponi 2016).

The T-type calcium channels display small single-channel conductance (T stands for transient or tiny) (Perez-Reyes et al. 1998) and are encoded by the CACNA1G (Ca_V_3.1), CACNA1H (Ca_V_3.2) and CACNA1I (Ca_V_3.3) genes. The Ca_V_3s have similar channel activation and inactivation kinetics and are ubiquitously expressed in the nervous, neuroendocrine, reproductive and cardiovascular systems (Perez-Reyes 2003, Hansen 2015). In heterologous expression systems, Ca_V_3.3 mediated currents display the slowest activation and inactivation kinetics and recover faster from inactivation (Kozlov et al. 1999), however due to this relatively small differences, they are nearly indistinguishable from the other T-type isoform in native tissue. In the brain, abundant CACNA1I transcripts display remarkable regional distribution and appear to prevail in distal dendrites (Perez-Reyes 2003, Lee et al. 1999), where currents mediated by Ca_V_3.3 mediate the major sleep spindle pacemaker in the thalamus (Astori et al. 2011). Transgenic mice lacking Ca_V_3.3 channels display severe alterations to sleep-spindle generator rhythmogenic properties posing this T-type channel isoform as a critical target in the study of brain function and development (Astori et al. 2011). However, the expression of Ca_V_3.3 channels overlaps with that of Ca_V_3.1 and/or Ca_V_3.2, which together with the lack of robust, isoform selective pharmacology has hampered the elucidation of their specific contributions to cellular physiology.

T-type calcium channels share the characteristic modular topology of other voltage-gated ion channels (VGIC) (Catterall et al. 2005) that consists of a voltage sensor (VS) module formed by transmembrane segments S1 through S4 and a pore module (PM) composed of the transmembrane segments S5 and S6 connected by a re-entrant pore loop. The VS controls channel opening in response to changes in membrane potential and the PM provides aqueous passage for ions across the lipid membrane. The tetrameric arrangement of four PMs lining the permeation pathway surrounded by four VSs in either swapped or non-swapped configuration enables VGIC function (for review see (Barros et al. 2019)). In voltage-gated potassium (K_**V**_) and transient receptor potential (TRP) channels, each monomer (1xVS + 1xPM) is encoded by a core α-subunit; whereas in voltage-gated sodium (Na_V_) and Ca_V_ channels the α-subunit contains the four homologous, but not identical, domains (DI-DIV) joined through large intracellular linkers (Catterall 2000, Catterall et al. 2005).

Natural compounds that evolved to occlude ion channel’s PM or to interact with their VS are distinguished broadly as pore blockers and gating modifiers, respectively. In VGICs, the extracellularly exposed areas of the VS are pharmacological targets of neurotoxins and synthetic compounds where at least three distinct pharmacological sites have been described in Na_V_ channels (Catterall et al. 2007). Within the VS is a conserved S3b-S4 paddle motif which can be transplanted into the VS of other VGICs and retain toxin sensitivity (Bosmans, Martin-Eauclaire, and Swartz 2008). α-Scorpion toxins, sea-anemone toxins and numerous spider toxins interact with Na_V_ channel site 3, located in the extracellular loop between DIV-S3 and DIV-S4 thereby interfering with the conformational changes that couple channel activation to fast inactivation (Hanck and Sheets 2007). Binding of β-scorpion toxins to the S1–S2 and S3–S4 loops of domain II (site 4) shifts the voltage dependence of channel activation towards depolarized voltages reducing the maximal current at normal activation potentials. Lastly, the pharmacological site 6 (located near site 3) is targeted by the δ-conotoxins that slow Na_V_ channel inactivation (Terlau et al. 1996). All these gating modifier peptides (GMPs) appear to “hold” the VS in different conformations leading to their mechanism of modulation to be recognized as “voltage-sensor trapping” (Cestele et al. 1998), a phenomenon also observed in the interaction of theraphotoxins with K_**V**_2.1 (Swartz and MacKinnon 1997) and agatoxins with Ca_V_ channels (McDonough, Mintz, and Bean 1997).

Sequence and functional conservation between the VSs leads to promiscuous interactions between peptides and small molecules across VGIC families. Examples of these include ProTx-I (Na_V_/K_**V**_/Ca_V_/TRPA1) (Bladen et al. 2014, Bosmans, Martin-Eauclaire, and Swartz 2008, Gui et al. 2014, Middleton et al. 2002); ProTx-II (Na_V_/Ca_V_) (Bladen et al. 2014, Middleton et al. 2002); Kurtoxin (Na_V_/Ca_V_) (Chuang et al. 1998); Hanatoxin (K_**V**_/Na_V_/Ca_V_) (Bosmans, Martin-Eauclaire, and Swartz 2008, Li-Smerin and Swartz 1998, Swartz and MacKinnon 1997), as well as JZTX-I (Na_V_/K_**V**_) (Xiao et al. 2005, Yuan et al. 2007). Furthermore, small molecules such as capsaicin, capsazepine (TRPV1/K_**V**_/Ca_V_) (Caterina et al. 1997, Kuenzi and Dale 1996, McArthur, Finol-Urdaneta, and Adams 2019), and A803467 (Na_V_, Ca_V_) (Bladen and Zamponi 2012) amongst others are known to interact across VGIC families.

Shared ancestry and sequence conservation within the voltage sensing machinery have been used to rationalize commonalities in structure, gating kinetics and pharmacophores between Na_V_ and T-type channels (Bladen and Zamponi 2012). Several Na_V_-active GMPs were shown to inhibit Ca_V_3.1 channels through interactions with the channel’s domain III (Ca_V_3.1^DIII^) (Salari et al. 2016); whereas the potent Na_V_1.7 inhibitor, μ-theraphotoxin Pn3a (Deuis et al. 2017a) also interacts with HVA Ca_V_ channels (McArthur et al. 2020).

In this study, we have used whole-cell patch clamp electrophysiology, mutagenesis and computational docking to probe Pn3a’s interactions with the LVA Ca_V_3 channels. Our results support the use of Pn3a as a molecular tool for the study of Ca_V_3.3-mediated currents in native cells and highlights a previously unrecognized pharmacophore that may enable selective targeting of T-type channel isoforms.

## Results

### Pn3a selectively inhibits Ca_V_3.3 channels

Functional assessment of Pn3a activity was examined on depolarization-activated calcium currents (I_Ca_) through the human T-type calcium channel isoforms: Ca_V_3.1, Ca_V_3.2 and Ca_V_3.3. Whole-cell currents were elicited by a 100 ms test pulse to −20 mV from a holding potential (Vh) of −90 mV at a frequency of 0.2 Hz and recorded at room temperature (20-22°C, Figure 1A). Pn3a (10 μM) strongly inhibited Ca_V_3.3 mediated currents (90.2 ± 1.9%, n = 5) with negligible effects over the two other T-type isoforms (Ca_V_3.1: 3.8 ± 1.8%, n = 5; Ca_V_3.2: 2.5 ± 2.5%, n = 5) (Figure 1A,B). These results indicate that Pn3a has >100-fold preference for Ca_V_3.3 over the other highly homologous Ca_V_3 isoforms. Scaled Ca_V_3.3 macroscopic currents recorded in the absence (control) and presence of Pn3a (3 μM) display similar macroscopic activation kinetics (τ_act, control_= 5.95 ± 0.54 ms vs τ_act, Pn3a_= 6.37 ± 0.44 ms; P =0.32, n = 5, paired t-test) while Pn3a slowed macroscopic inactivation kinetics (τ_inact control_= 39.91 ± 8.25 ms vs τ_inact, Pn3a_= 60.25 ± 16.02 ms; P = 0.04, n = 5, paired t-test) (representative currents provided in the inset, Figure 1A). The preferential actions of Pn3a highlight its potential as a molecular probe to isolate and study the contribution of Ca_V_3.3 currents in native cells.

**Figure 1.**
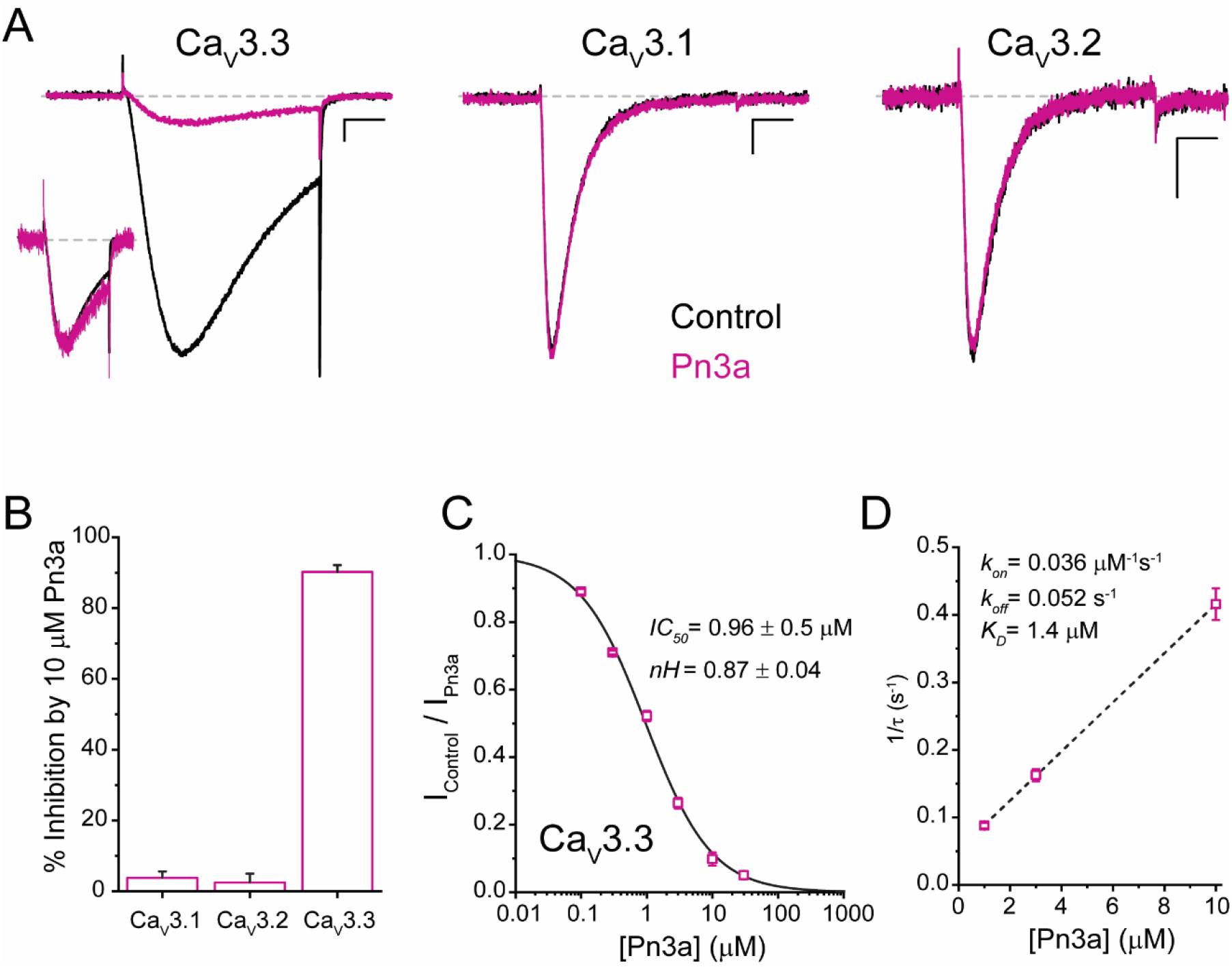
Pn3a preferentially inhibits human Ca_V_3.3-mediated Ca^2+^ currents. **(A)** Ca_V_3.3 (left), Ca_V_3.1 (middle), and Ca_V_3.2 (right) currents elicited by 100 ms step depolarization to -20 mV (Vh - 90 mV, 0.2 Hz) in the absence (control, black) and presence of 3 μM Pn3a (pink). Scale bars: 0.2 nA, 20 ms. The inset shows scaled Ca_V_3.3 currents in control and in the presence of Pn3a with similar kinetics. **(B)** Bar graph summarizing % inhibition by 10 μM Pn3a of the three Ca_V_3 isoforms. **(C)** Concentration-response curve for Pn3a inhibition of Ca_V_3.3 currents. **(D)** Kinetics of Pn3a inhibition of Ca_V_3.3. K_obs_ was determined at three concentrations and fit with a linear equation where K_obs_ = k_on_ · [Pn3a] + k_off_.

Pn3a inhibitory potency against Ca_V_3.3 currents was assessed at increasing peptide concentrations from which a concentration-response curve was built (Figure 1C). Fit to a standard Hill equation rendered an *IC*_*50*_ value equal to 0.96 ± 0.05 μM (*nH* 0.87 ± 0.04, n = 5 per concentration) for the inhibition of Ca_V_3.3 channels. This Hill coefficient is consistent with a 1:1 stoichiometry between Pn3a toxin and Ca_V_3.3 channels.

The change in Ca_V_3.3 peak current amplitude during Pn3a *washin* and *washout* enables the assessment of Pn3a binding to Ca_V_3.3 channels. The inhibition of Ca_V_3.3 calcium currents by Pn3a was mono-exponential with progressively faster tau values (τ_obs_) at increasing peptide concentration (Figure 1D). The on- and off-rate constants (*k*_*on*_ and *k*_*off*_) were determined from the fit to the linear plot of 1/τ_obs_ versus [Pn3a] (Figure 1D, n = 5 per concentration), where *k*_*on*_ is the slope and *k*_*off*_ is the y-intercept. The linear regression line results in a *k*_*on*_ of 0.036 μM^-1^sec^-1^ and *k*_*off*_ of 0.052 sec^-1^ which yield a *K*_*D*_ of 1.4 μM, in close agreement with the IC_50_ value obtained (Figure 1C). *k*_*off*_ was confirmed by fitting the *washout* to a single exponential (*k*_*off*_ = 0.055 ± 0.002 sec^-1^). Hence, Pn3a inhibits Ca_V_3.3 mediated currents without apparent actions on Ca_V_3.1 or Ca_V_3.2 exposed to up to 10 μM peptide.

### Pn3a modifies Ca_V_3.3’s gating

The voltage dependence of Ca_V_3.3 activation, deactivation, inactivation and recovery from inactivation were investigated in the absence and presence of Pn3a (3 μM) (Figure 2 and Table 1). Ca_V_3.3 activation is shifted ∼13 mV to more depolarized potentials (control V_0.5_= -28.3 ± 0.4 mV, n = 5, vs Pn3a V_0.5_ = -15.3 ± 0.4 mV, n = 5; P<0.0001), indicating that a stronger depolarization is required to enable channel opening in the presence of Pn3a (Figure 2A,C). The voltage dependence of Ca_V_3.3 steady-state inactivation (SSI) was not affected by exposure to the spider peptide (control V_0.5_= -56.3 ± 0.3 mV, n = 5, vs Pn3a V_0.5_ = -56.6 ± 0.4 mV, n = 5, P = 0.57, Figure 2B-C), yet Ca_V_3.3-mediated currents inactivated by a 200 ms pre-pulse to –20 mV recovered faster in the presence of Pn3a (τ = 0.22 ± 0.01 s, n = 5) than under control conditions (τ = 0.34 ± 0.01 s, n = 5; P < 0.0001) (Figure 2D-E).

**Figure 2.**
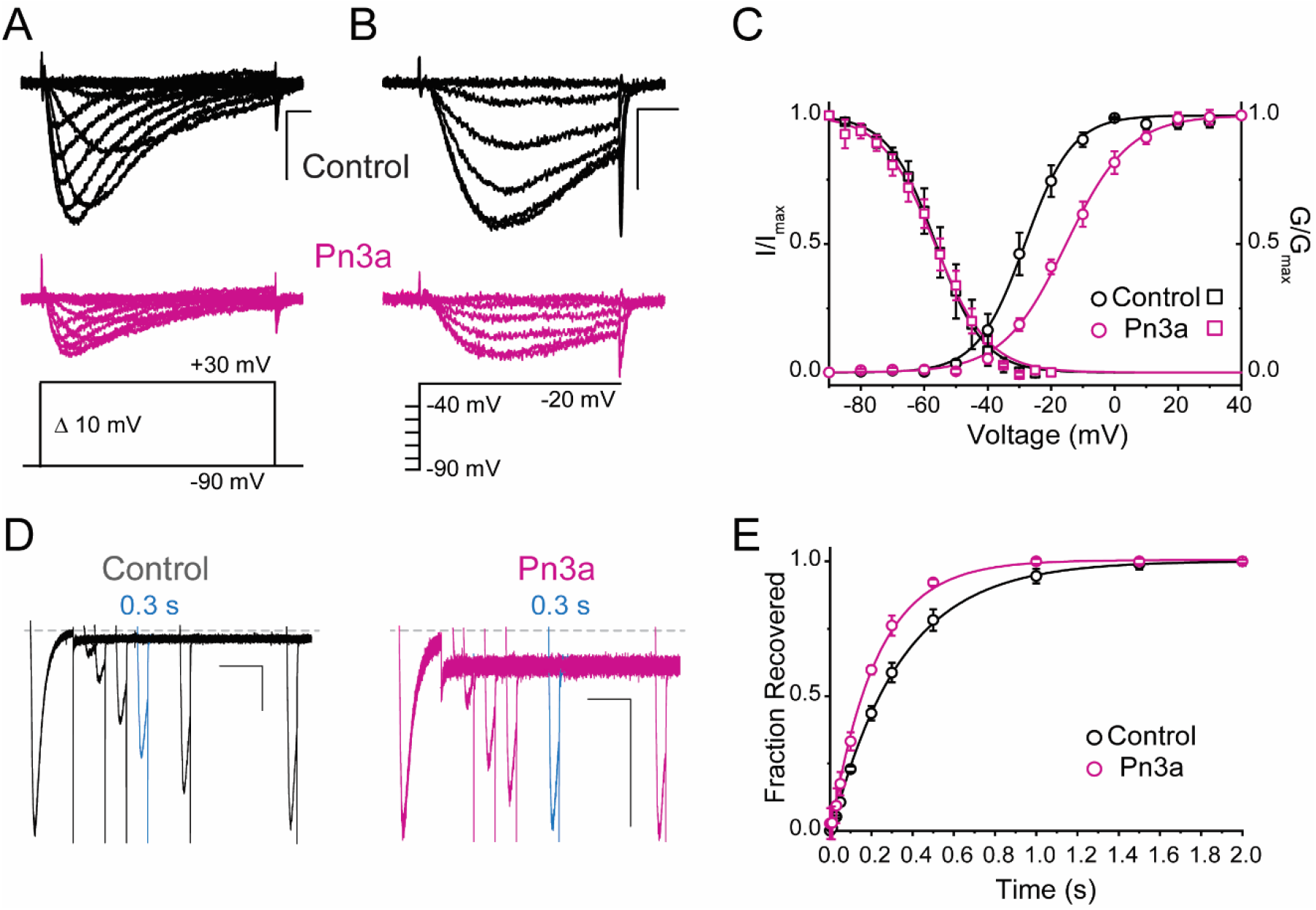
Pn3a produces a depolarizing shift in the voltage dependence of activation and speeds up recovery from inactivation of Ca_V_3.3. **(A-C)** Effect of Pn3a on the voltage dependence of Ca_V_3.3. Representative currents from **(A)** activation or **(B)** steady-state inactivation protocols, in the absence (Top: Control, Black) and presence of 3 μM Pn3a (middle: Pn3a, pink) using the standard protocols (bottom). Scale bars: 0.5 nA, 10 ms. **(C)** Activation (circles) and steady-state inactivation (squares) relationships for Ca_V_3.3 in the absence (black) and presence of 3 μM Pn3a (pink). **(D)** Representative recovery from inactivation currents in control (left) and presence of 3 μM Pn3a (right) (trace shown in blue highlights the current recovered after 0.3 s). Scale bars: 0.2 nA, 200 ms. **(E)** Recovery from inactivation in the absence (black) and presence of 3 μM Pn3a (pink).

**Table 1.** Activation and inactivation values in control and presence of Pn3a (3 μM for hCa_V_3.3; 1 μM for K_**V**_1.7-Ca_V_3.3^D1-IV^ chimeras). * Significant determined from paired t-test with significance threshold set to P < 0.05.

|  |  | Control | Pn3a |
| --- | --- | --- | --- |
| hCav3.3 | *V*<sub>0.5</sub> Activation | -28.3 ± 0.4 mV (5) | -15.3 ± 0.4 mV (5) * |
|  | *k*<sub>a</sub> Slope factor | 7.5 ± 0.3 (5) | 10.0 ± 0.3 (5) * |
|  | *V*<sub>0.5</sub> SSI | -56.3 ± 0.3 mV (5) | -56.6 ± 0.4 mV (5) |
|  | *k*<sub>a</sub> Slope factor | 7.5 ± 0.2 (5) | 8.4 ± 0.4 (5) |
|  | τ Recovery | 0.22 ± 0.01 ms (5) | 0.34 ± 0.01 (5) * |
| Kv1.7 | *V*<sub>0.5</sub> Activation | -10.5 ± 0.6 mV (7) | - |
|  | *k*<sub>a</sub> Slope factor | 9.9 ± 0.5 (7) | - |
| Kv1.7-Cav3.3<sup>DI</sup> | *V*<sub>0.5</sub> Activation | 91.3 ± 0.4 mV (5) | 88.2 ± 0.3 mV (5) |
|  | *k*<sub>a</sub> Slope factor | 1.0 ± 0.3 (5) | 11.8 ± 0.2 (5) |
| Kv1.7-Cav3.3<sup>DII</sup> | *V*<sub>0.5</sub> Activation | 97.3 ± 0.4 mV (9) | 112.6 ± 0.4 mV (7) * |
|  | *k*<sub>a</sub> Slope factor | 14.1 ± 0.3 (9) | 12.3 ± 0.4 (7) * |
| Kv1.7-Cav3.3<sup>DIII</sup> | *V*<sub>0.5</sub> Activation | 71.3 ± 0.8 mV (7) | 74.7 ± 0.9 mV (5) |
|  | *k*<sub>a</sub> Slope factor | 21.1 ± 0.7 (7) | 22.0 ± 0.8 (5) |
| Kv1.7-Cav3.3<sup>DIV</sup> | *V*<sub>0.5</sub> Activation | -60.8 ± 0.8 mV (5) | -60.8 ± 0.4 mV (5) |
|  | *k*<sub>a</sub> Slope factor | 10.1 ± 0.7 (5) | 10.0 ± 0.4 (5) |

The voltage-dependence of Pn3a inhibition of I_Ca_ and Na^+^ mediated (I_Na_) Ca_V_3.3 currents was examined at various test potentials (Figure 3A and Figure 3-figure supplement 1). Pn3A displayed a decrease in Ca_V_3.3 I_Ca_ inhibition at more depolarized potentials consistent with Pn3a preferentially binding to the down-state of the voltage sensor. To examine the voltage dependence across a larger voltage range, extracellular Ca^2+^ was removed permitting Na^+^ to be the primary charge carrier through the open Ca_V_3.3. Similar to that seen for I_Ca_, I_Na_ showed a similar voltage dependence, with stronger inhibition at more negative potentials (Figure 3-figure supplement 1).

**Figure 3.**
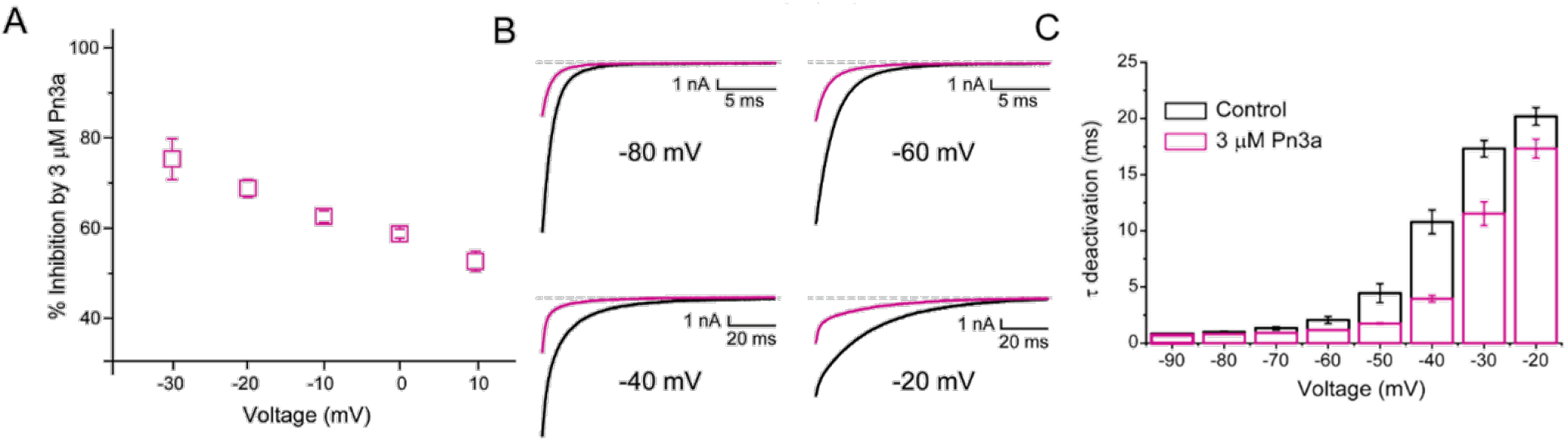
Pn3a inhibition of Ca_V_3.3 is stronger at hyperpolarized potentials and speeds up channel deactivation. **(A)** Voltage dependence of Pn3a inhibition of Ca_V_3.3 mediated Ca^2+^ currents. **(B)** Representative Ca_V_3.3 tail currents at different voltages in the absence (black) and presence (pink) of 3 μM Pn3a. **(C)** Summary of the time constant (τ) of Ca_V_3.3 deactivation upon return to the holding potential (−90 mV) plotted against the activating (pre-pulse) potential in control (black) and in the presence of Pn3a (3 μM, pink).

The time course of tail current decay reflects the rate of channels leaving the open state (test potential) and entering the closed state (deactivation) upon return to the holding potential. We examined the modulation of Ca_V_3.3 channel deactivation through tail current kinetic analysis. Pn3a-modified Ca_V_3.3 currents displayed faster deactivation *tau* values (τ_deactivation_) across all potentials tested compared to control (Figure 3 B-C). In combination with the observed increase in recovery from inactivation, the data suggests that Pn3a acts via destabilizing the open/inactivated state or stabilizing the closed state. These results suggest that Pn3a is a Ca_V_3.3 gating modifier peptide that decreases channel availability by increasing the energy required for channel opening.

## Control Pn3a

### Pn3a interacts with Ca_V_3.3’s DII S3-S4 paddles

To ascertain Pn3a’s pharmacophore on Ca_V_3.3 channels, we applied the chimeric approach of transplanting its four S3-S4 paddles into a voltage-gated potassium channel based on a similar template to that previously described for K_**V**_2.1/Ca_V_3.1 chimeras (Salari et al. 2016). We substituted portions of the S3b-S4 from Ca_V_3.3 DI to DIV into the K_**V**_1.7 channel backbone. The sequence alignment of Ca_V_3.3 DI-DIV S3-S4 segments and corresponding extracellular region of K_**V**_1.7 is presented in Figure 4A.

**Figure 4.**
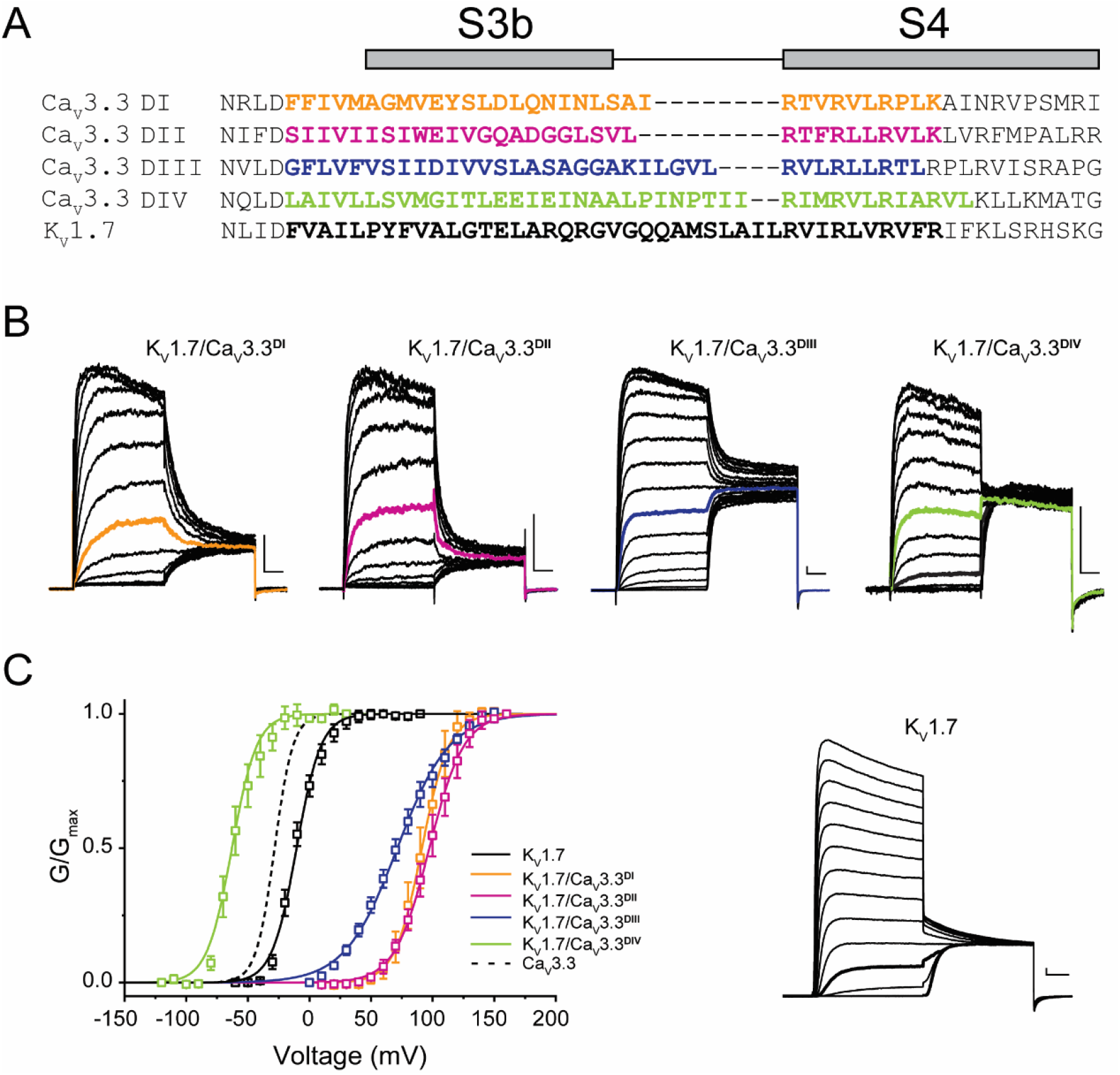
Chimeric constructs Ca_V_3.3 voltage sensor paddles and the K_V_1.7 channel. **(A)** Sequence alignment between paddle regions of Ca_V_3.3 and K_**V**_1.7. The coloured bolded sequences from Ca_V_3.3 DI (yellow), Ca_V_3.3 DII (pink), Ca_V_3.3 DIII (blue), Ca_V_3.3 DIV (green) were grafted onto K_**V**_1.7 (black). **(B)** Representative current traces in response to 50 ms long I-V protocols used to evaluate the voltage dependence of activation of all constructs (Vh = -80 mV). K_**V**_1.7/Ca_V_3.3^DI^ (0 to 160 mV), K_**V**_1.7/Ca_V_3.3^DII^ (0 to 160 mV), K_**V**_1.7/Ca_V_3.3^DIII^ (0 to 160 mV) and K_**V**_1.7/Ca_V_3.3^DIV^ (−120 to 30 mV). The traces highlighted in colour correspond to currents near half-activation potential (V_0.5_). **(C)** Activation curves for all chimeras: K_**V**_1.7/Ca_V_3.3^DI^ (yellow), K_**V**_1.7/Ca_V_3.3^DII^ (pink), K_**V**_1.7/Ca_V_3.3^DIII^ (blue) and K_**V**_1.7/Ca_V_3.3^DIV^ (green) and the parental channel K_**V**_1.7 (black). The dotted line corresponds to Ca_V_3.3 activation for reference. The inset contains representative K_**V**_1.7 currents (−60 to 90 mV). All scale bars: 1 nA, 10 ms.

All four K_**V**_1.7/Ca_V_3.3 chimeric constructs were functional, mediating large K^+^ currents that displayed distinct gating properties from those of the parental wild-type K_**V**_1.7 (V_0.5_ = -10.5 ± 0.6 mV, n = 7; Figure 4B, inset). Briefly, chimeric K_**V**_1.7/Ca_V_3.3^DI^, K_**V**_1.7/Ca_V_3.3^DII^ and K_**V**_1.7/Ca_V_3.3^DIII^ displayed 80-100 mV depolarizing shifts in channel activation (K_**V**_1.7/Ca_V_3.3^DI^ V_0.5_ = 91.3 ± 0.4 mV, n = 5; K_**V**_1.7/Ca_V_3.3^DII^ V_0.5_ = 97.3 ± 0.4 mV, n = 9; K_**V**_1.7/Ca_V_3.3^DIII^ V_0.5_ = 71.3 ± 0.8 mV, n = 7, two-way ANOVA, P < 0.05), whereas K_**V**_1.7/Ca_V_3.3^DIV^ activation was ∼50 mV hyperpolarized (K_**V**_1.7/Ca_V_3.3^DIV^ V_0.5_ = -60.8 ± 0.8 mV, n = 5; two-way ANOVA, P < 0.05) (Figure 4). Our results are in broad agreement with previous reports for Ca_V_3.1, where DIV displays a hyperpolarizing shift and DI-III show a variable depolarizing shift in V_0.5_ (Salari et al. 2016).

K_**V**_1.7/Ca_V_3.3^DI-IV^ chimeras and the parental K_**V**_1.7 construct were analysed in control and after exposure to Pn3a (1 μM) and representative currents (Vh = -100 mV, with test pulse to +40 mV (K_**V**_1.7), +120 mV (K_**V**_1.7/Ca_V_3.3^DI-III^) or 0 mV (K_**V**_1.7/Ca_V_3.3^DIV^)) in both conditions are included as insets (Fig 5). Similar to the parental channel, peak currents of K_**V**_1.7/Ca_V_3.3^DI^ and K_**V**_1.7/Ca_V_3.3^DIV^ chimaeras were insensitive to Pn3a (K_**V**_1.7: 1.0 ± 0.8%, n = 6; Ca_V_3.3^DI^: 0.6 ± 0.5%, n = 5, and Ca_V_3.3^DIV^: 0.4 ± 0.2%, n = 5) (Figure 5A, D, E), with K_**V**_1.7/Ca_V_3.3^DIII^ displaying modest peptide-dependent inhibition (12.0 ± 0.6%, n = 6) (Figure 5C).

**Figure 5.**
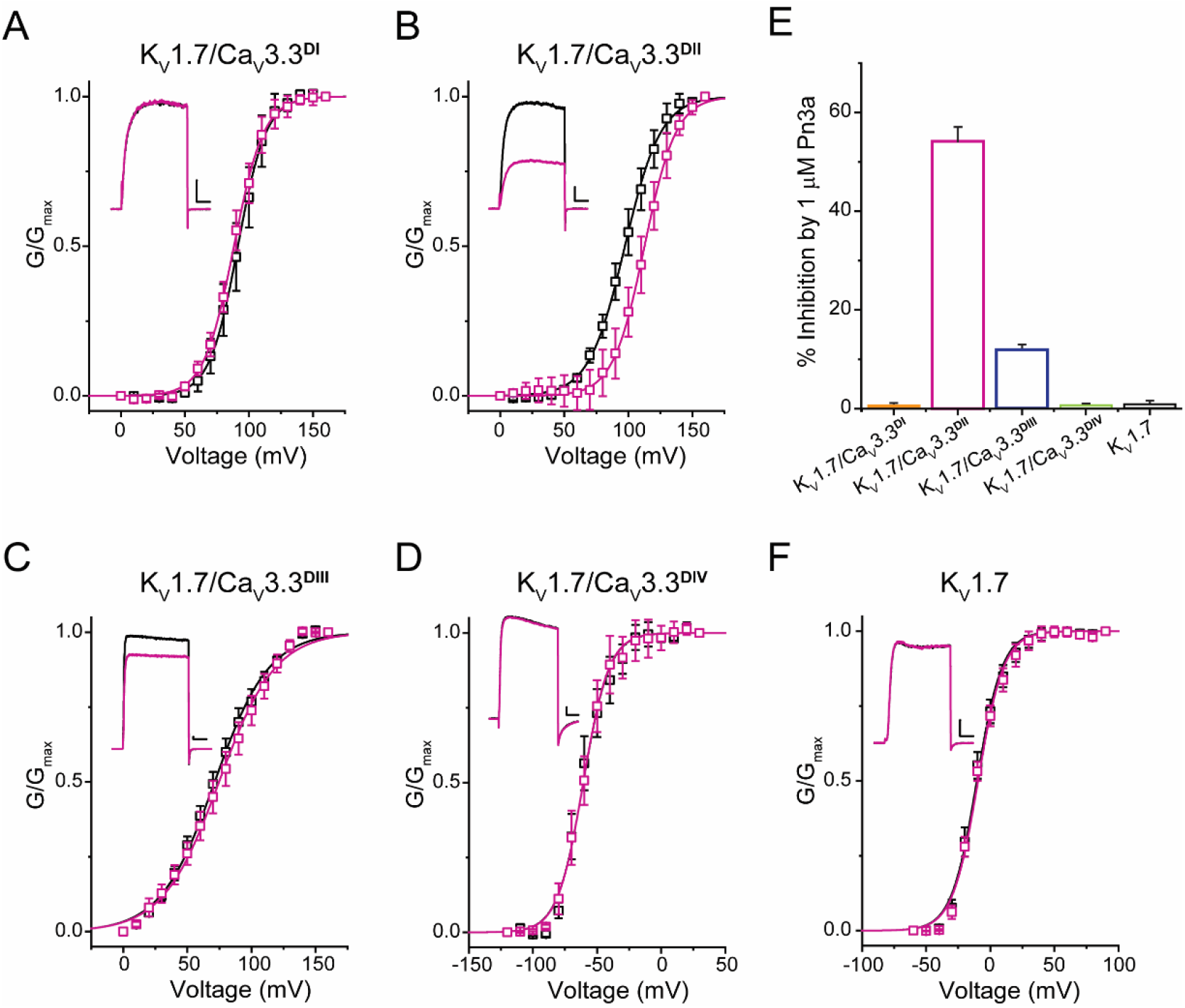
Pn3a shifts the activation of K_V_1.7/Ca_V_3.3^DII^ chimeric channels. G/G_max_-V relationships and representative currents obtained in the absence (control, black) and presence of 1 μM Pn3a (pink) for **(A)** K_**V**_1.7/Ca_V_3.3^DI^ (90 mV), **(B)** K_**V**_1.7/Ca_V_3.3^DII^ (100 mV), **(C)** K_**V**_1.7/Ca_V_3.3^DIII^ (70 mV), and **(D)** K_**V**_1.7/Ca_V_3.3^DIV^ (−60 mV). **(E)** Bar graph showing percent inhibition by 1 μM Pn3a of K_**V**_1.7/Ca_V_3.3^DI-IV^ chimeras and K_**V**_1.7. **(F)** K_**V**_1.7 G/G_max_-V relationship and current traces obtained in the absence (control) and presence of Pn3a (−10 mV).

In contrast, currents mediated by K_**V**_1.7/Ca_V_3.3^DII^ were significantly reduced (54.2 ± 2.9%, n = 7) in the presence of Pn3a (Figure 5B). Furthermore, conductance-voltage relationships for each S3-S4 paddle chimera in the absence and presence of Pn3a were built from peak currents in response to standard I-V protocols (K_**V**_1.7 -60 to +90 mV; K_**V**_1.7/Ca_V_3.3^DI-III^ 0 to +160 mV; and K_**V**_1.7/Ca_V_3.3^DIV^ -120 to +30 mV; Vh = -100 mV, 0.2 Hz). The latter analysis revealed a ∼15 mV rightward shift in the half voltage of activation (V_0.5_) exclusively in the Ca_V_3.3^DII^ chimera when exposed to Pn3a (control V_0.5_ = 97.3 ± 0.4 mV, n = 9; vs Pn3a V_0.5_ = 112.6 ± 0.4 mV, n = 7, p < 0.0001), strongly suggesting that Pn3a interacts with the S3-S4 region of domain II in Ca_V_3.3. These results suggest that the positive shift in activation V_0.5_ observed in the full length and DII chimera likely underpins Pn3a’s mechanism of Ca_V_3.3 inhibition, highlighting the differences in pharmacological properties of each voltage-sensing module.

### Molecular determinants of Pn3a inhibition of Ca_V_3.3 over other Ca_V_3 isoforms

To predict the molecular determinants responsible for the selective inhibition of Ca_V_3.3 over other Ca_V_3 isoforms by Pn3a, molecular docking was used to assess the most energetically favoured binding poses for Pn3a on each of the individual VS of the three channel isoforms, Ca_V_3.3, Ca_V_3.2, and Ca_V_3.1 (Figure 6 and Figure 6-figure supplement 1).

**Figure 6.**
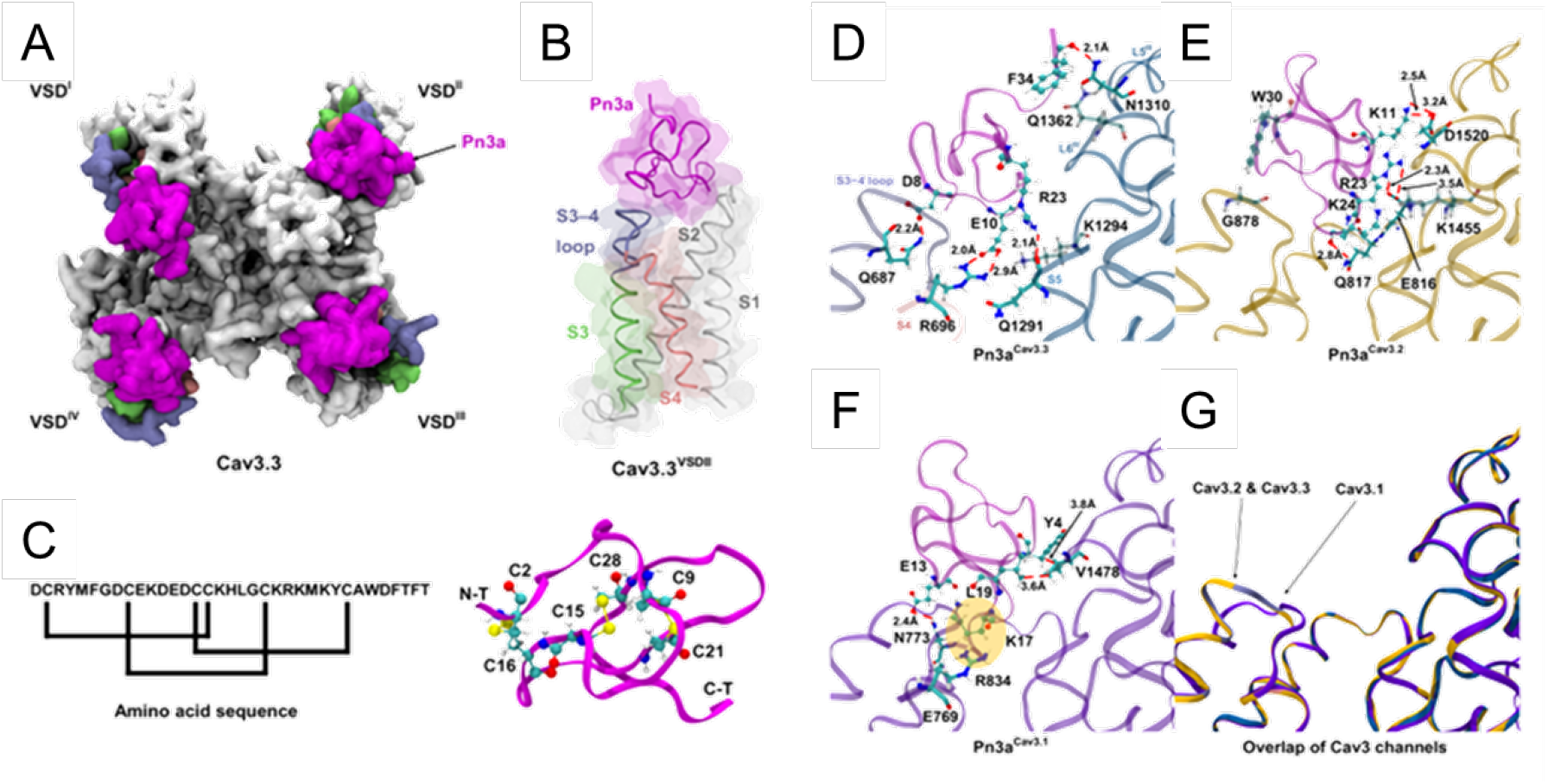
Pn3a/Ca_V_3 binding conformations. The homology models of Ca_V_3.3 and Ca_V_3.2 were built after the Cryo-EM structure of Ca_V_3.1 (PDB ID: 6KZO (Zhao et al. 2019)) using SWISS-MODEL (Arnold et al. 2006). (**A**) Top view of Pn3a (magenta) binding to the four domain voltage sensors of Ca_V_3.3. The S3 (lime), S3-4 loop (ice blue) and S4 (pink) are highlighted. (**B**) Side view of the binding conformation of Pn3a to Ca_V_3.3^DII^. (**C**) Left: Pn3a amino acid sequence and disulfide connectivity. Right: 3-D structure of Pn3a (PDB ID: 5T4R (Deuis et al. 2017b)) showing cysteine residues in CPK linked by salt-bridges (yellow). Pairwise amino acid interactions between Pn3a (magenta) with the extracellular loop of: **(D)** Ca_V_3.3^DII^ (dark blue), **(E)** Ca_V_3.2^DII^ (yellow), and **(F)** Ca_V_3.1^DII^ (purple). **(G)** Superposition between the Ca_V_3.1 Cryo-EM structure and the homology models of Ca_V_3.3 and Ca_V_3.2 highlighting the difference the extracellular S3-S4 loop between Ca_V_3.3 and the other two Ca_V_3 isoform. **(D-G)** S1, S1-2 loop and S2 were removed for clarity.

Comparative analyses of toxin interactions with the other T-type calcium channel members showed that Pn3a forms fewer contacts with the DII S3-S4 linker of Ca_V_3.1 and Ca_V_3.2, compared to Ca_V_3.3 (Figure 6 E-G). For Ca_V_3.2, the orientation of bound Pn3a shows a similar orientation towards the S1-S2/S3-S4 linker Ca_V_ity as that at Ca_V_3.3 (Figure 6E/G). However, the pairwise interactions formed by Pn3a’s residues K11, R23 and K24 and those on the S1-S2 and extracellular region L6III of Ca_V_3.2 (specifically with D1520, E816 and Q817), render loops 2 and 4 of Pn3a to be shifted closer to DIII’s extracellular regions and S1-S2 loop (Figures 6E/G), respectively, rather than towards the S3-S4 loop and S5 as seen for Ca_V_3.3. As a consequence, the cationic residues on loops 1 and 2 of Pn3a established fewer contacts with Ca_V_3.2^DII^’s S3, S4 and S3-4 linker.

In the case of Ca_V_3.1, hydrophobic interactions formed by Pn3a-Y4 and L19 with Ca_V_3.1^DIII^’s S6 extracellular loop (V1478), accompanied by hydrogen bonding between E13 and N773 (VS DII-S2), render the peptide loop 1 in proximity to the pore loop of Ca_V_3.1^DIII^ (Figure 6F/G and Figure 6-figure supplement 1). These interactions also position loop 3 of the peptide in the cavity between the S1-S2 and S3-4 linkers of Ca_V_3.1. The electrostatic clash imposed by this binding position between Pn3a-K17 (loop 3) and Ca_V_3.1^DIII^ S4’s 2nd gating charge (R834), as shown in Figure 6F, may contribute to Ca_V_3.1’s insensitivity to Pn3a modulation.

## Discussion

The present study demonstrates the functional activity of Pn3a as a gating modifier inhibitor of human Ca_V_3.3 channels with >100-fold higher activity over Ca_V_3.1 and Ca_V_3.2. Our chimeric approaches revealed Pn3a’s preference for Ca_V_3.3^DII^ S3-S4 paddle region, whereas comparative molecular docking amongst isoforms identified a novel binding site putatively determining Pn3a’s proclivity towards Ca_V_3.3. Thus, this investigation highlights Pn3a as the first molecular probe available for the study of Ca_V_3.3 contribution in native cells. We propose that the unique features of Pn3a’s Ca_V_3.3-specificity may be exploited to design isoform-selective T-type calcium channel modulators.

### Ca_V_3 sub-type selectivity

μ-Theraphotoxin-Pn3a, was originally described as a highly specific Na_V_1.7 inhibitor (Deuis et al. 2017a) and more recently shown also to inhibit HVA calcium channels (McArthur et al. 2020). Perhaps a more interesting aspect of Pn3a is its >100-fold selectivity for human Ca_V_3.3 (IC_50_ of 960 nM) over both Ca_V_3.1 and Ca_V_3.2 (Figure 1) which distinguished this as a unique isoform-selective compound against the T-type calcium channels. Only a handful of venom-derived peptides have been shown to interact with T-type calcium channels. The scorpion peptide Kurtoxin was the first GMP reported to modulate Ca_V_3 channels with high affinity for Ca_V_3.1 and Ca_V_3.2 channels (Chuang et al. 1998). The closely related peptide KLI, from *Parabuthus granulatus*, was later shown to inhibit Ca_V_3.3 with and IC_50_ ∼450 nM (Olamendi-Portugal et al. 2002). However, the T-type isoform selectivity and binding site of these two peptides was not comprehensively documented at the time. The activity of spider Protoxins I and II against the three Ca_V_3 channels revealed that ProTx-I preferentially modulates Ca_V_3.1 channels whereas ProTx-II targets Ca_V_3.2 (Bladen et al. 2014, Salari et al. 2016). Thus, Pn3a complements the molecular toolbox for the study of T-type calcium channels in native tissues.

### Voltage dependence and current kinetics

Pn3a inhibits Ca_V_3.3 by inducing a depolarizing shift in Ca_V_3.3 voltage dependence of activation in a manner analogous to Kurtoxin, Protoxin I and Protoxin II actions on Ca_V_3.1 channels (Edgerton, Blumenthal, and Hanck 2010, Chuang et al. 1998). This is also consistent with Pn3a’s modulation of Na_V_1.7 (Deuis et al. 2017a) but not of Ca_V_2.2 channels for which a hyperpolarizing shift in the voltage dependence of inactivation was associated with current inhibition (McArthur et al. 2020). In contrast to the 2-fold slower recovery from inactivation reported for Pn3a-bound Na_V_1.7 channels, currents mediated by Pn3a-modified Ca_V_3.3 currents recover ∼1.5-fold faster from inactivation (Figure 2D/E) indicative of a toxin-induced destabilization of the inactivated state.

Pn3a inhibition of Ca_V_3.3 was voltage dependent with greater inhibition at more negative potentials (Figure 3A) similar to Kurtoxin-bound Ca_V_3.1 channel currents (Chuang et al. 1998). However, a delay in Ca_V_3.1 activation kinetics was apparent in the presence of both Kurtoxin and ProTx-II (Edgerton, Blumenthal, and Hanck 2010, Chuang et al. 1998) but not in Pn3a-modified Ca_V_3.3. The spider peptides Pn3a (this study) and ProTx-II (Edgerton, Blumenthal, and Hanck 2010) appear to slow T-type channel deactivation suggestive of a stabilization of the channel’s closed state, whereas Kurtoxin, from scorpion venom, does not affect Ca_V_3.1 channel closure (Chuang et al. 1998) highlighting incompletely understood aspects of GMP/VGIC interactions.

### Interaction of Pn3a with Ca_V_3.3 VS paddles

The portability of the S3-S4 paddle region was shown more than 20 years ago (Swartz and MacKinnon 1997). To date most studies, if not all, have used the K_**V**_2.1 channel backbone for the identification of GMPs binding sites in voltage-gated sodium and calcium channels expressed in *Xenopus* oocytes (Bosmans, Martin-Eauclaire, and Swartz 2008, Salari et al. 2016). For our chimera studies, we selected the K_V_1.7 backbone given the remarkable scarcity of GMPs interacting with K_**V**_1 channels (Finol-Urdaneta et al. 2020). The robust expression of this channel in heterologous systems (Finol-Urdaneta, Struver, and Terlau 2006, Finol-Urdaneta et al. 2012) and resistance to Pn3a modulation (Figure 5E/F) make it suitable to assess peptide binding to Ca_V_3.3 paddle motifs in mammalian cells. The generated K_**V**_1.7/Ca_V_3.3 constructs resulted in four functional, voltage-gated chimeric potassium channels that exhibited distinct gating properties to those of the parental scaffold reflecting the acquisition of the grafted S3-S4 paddle regions from DI-DIV of Ca_V_3.3. Namely, K_**V**_1.7/Ca_V_3.3^DI-DIII^ all displayed >80 mV depolarizing shifts in channel activation compared to wild-type K_**V**_1.7, whereas the Ca_V_3.3^DIV^ bearing chimeric construct presented a comparable magnitude shift in the hyperpolarizing direction (Fig 4).

Analogous chimeric approaches of K_**V**_2.1/Ca_V_3.1D^I-IV^ have indicated that ProTx-II, PaTx-1, GsAF-I and GsAF-II exert their inhibitory actions predominantly through interaction with Ca_V_3.1^DIII^ (Salari et al. 2016), whereas Pn3a does so by targeting K_**V**_2.1/Na_V_1.7^DII^ and K_**V**_2.1/Na_V_1.7^DIV^ (Deuis et al. 2017a). Here we observe modest Pn3a inhibition of K_**V**_1.7/Ca_V_3.3^DIII^ mediated currents and potent effects on K_**V**_1.7/Ca_V_3.3^DII^ chimeras with current inhibition coupled to toxin-induced rightward shift in the voltage dependence of activation as evidence of preferential interactions with Ca_V_3.3’s DII (Figure 5B/F). A substantial body of literature has shown that DI-DIII and DIV are important for Na_V_ activation and inactivation, respectively (41-43). Thus, the predominant effects of Pn3a on Ca_V_3.3 activation are consistent with its interaction with DII of this channel.

Furthermore, it has been shown that ProTx-I inhibits Ca_V_3.3/Ca_V_3.1^DIV^ chimeric channels while interacting less potently with Ca_V_3.3/Ca_V_3.1^DII^. However, a clear binding site could not be delineated through mutation of individual Ca_V_3.1^DII^ residues as those did not result in measurable changes in toxin affinity (Bladen et al. 2014). It can be surmised that subtle, but significant GMP/VGIC interaction differences highlight incompletely understood idiosyncrasies related to molecular aspects of ion channel function as well as how peptide interactions may be affected by the experimental manipulation and conditions used for their study.

### A novel binding site with Ca_V_3 sub-type selectivity

Our molecular docking calculations suggest that Pn3a binds within the groove formed between extracellular linkers S1-S2 and S3-S4 supported by electrostatic interactions with the Ca_V_3.3^DII^ S3-S4 paddle, which differs substantially from the Ca_V_3.1/ProTx-II complex in which favourable binding sites appear in Ca_V_3.1’s DIV and DII (Bladen et al. 2014).

The interaction of Pn3a with Ca_V_3.3 channels involves substantial charge-charge interactions within the S3-S4 paddle (D8-Q687) and S4 (E10-R696) of Ca_V_3.3^DII^ (Figure 6). Moreover, favourable contacts between Pn3a (E10, R23 and F34) and Ca_V_3.3^DIII^’s S5 (Q1291), S5-S6 loop (K1294 and N1310) and S6 (Q1362) tilt the Pn3a C-terminus towards the pore-loop of DIII. This toxin pose is analogous to Dc1a’s insertion into the cleft between Na_V_PasD^II^ and the pore of the TTX-bound channel evidenced by Cryo-EM (Shen et al. 2018), as well as other previously reported Na_V_ channel GMPs (Katz et al. 2021). Conversely, the proximity between anionic residues (D8 and E10) on loops 1 and 2 of Pn3a and the Ca_V_3.3D^II^ S3-S4 paddle places this peptide at the hollow between S1-S2 and S3-S4 linkers, thus stablishing a binding conformation dissimilar from other tarantula toxins targeting Na_V_ channel site 4, like HwTX-IV, ProTx-II and HNTX-III, that bear critical basic and hydrophobic amino acids within loop 4 that interact with acidic residues on the Na_VDII_‘s S3-S4 linker (Deng et al. 2013, Liu et al. 2013, Bosmans and Swartz 2010). As opposed to SGTx1 and ProTx-II (Wang et al. 2004, Smith et al. 2007), the unique binding conformation of Pn3a to Ca_V_3.3 is guided by loop 2 anionic amino acid (D8, E10, D12, E13 and D14) interactions that position Pn3a within pharmacological site 4 of this T-type calcium channel, with overall lower contribution from basic peptide residues (Figure 6-figure supplement 1 and 2). Hence the predicted Pn3a interaction with Ca_V_3.3 is in contrast to that of previously identified spider peptide toxins modulating Na_V_ and K_**V**_ channels in which the common bioactive surface consists of positively charged and hydrophobic residues (Smith et al. 2007, Corzo et al. 2005).

Consistent with previous studies of tarantula toxins interactions with Na_V_ and K_**V**_ channels (Deng et al. 2013, Smith et al. 2007), our model signals important polar interactions between Pn3a and the three T-type calcium channels. However, in comparison to the multiple contacts observed in the Ca_V_3.3^DII^/Pn3a complexes, the homologous domains, Ca_V_3.1^DII^ and Ca_V_3.2^DII^ appear to form fewer interactions with Pn3a, consistent with our functional data. At the atomic level, the lack of inhibition of Ca_V_3.1 and Ca_V_3.2 mediated currents by Pn3a can be explained through ligand-receptor interactions. The basic residues of Pn3a, specifically K11, R23 and K24, constitute a ring that contacts Ca_V_3.2 polar and anionic residues D1520, E816 and Q817 via salt-bridges and hydrogen bonds (Figure 6E), similar to the interactions between Pn3a (D8, E10, R23 and F34) and Ca_V_3.3 (as mentioned above). However, Pn3a interactions with the Ca_V_3.2^DII^ VS were much fewer than those seen for Ca_V_3.3^DII^; whilst in the Ca_V_3.1D^II^/Pn3a bound complex, an electrostatic clash between R834 (Ca_V_3.1^DII^ S4’s 2^nd^ gating charge) and Pn3a-K17 (Figure 6F) poses a high energy barrier for the interactions between the toxin and Ca_V_3.1. In contrast, the homologous gating charge (R696) in Ca_V_3.3 is seen forming a favourable salt-bridge with Pn3a-E10. As VS gating charge displacement is required for the activation of voltage dependent ion channels (Lacroix and Bezanilla 2011), the E10-R696 interaction observed in the Ca_V_3.3^DII^/Pn3a docking complex may serve a critical role enabling Pn3a inhibition of Ca_V_3.3 mediated currents.

The findings presented here establish Pn3a as a gating modifier modulator of Ca_V_3.3 channels interacting with the paddle motif of DII’s voltage sensor module through putative stabilization of the closed/resting state and concomitant channel current inhibition. Pn3a’s >100-fold higher potency against Ca_V_3.3 over the other two Ca_V_3 isoforms is rationalized through recognition of a previously unknown drug binding site that may be exploited in the design of isoform-selective Ca_V_3 channel modulators.

## Materials and Methods

### Cell culture and transfections

Human embryonic kidney (HEK293T) cells containing the SV40 Large T-antigen were cultured and transfected by calcium phosphate method as reported previously (McArthur et al. 2018). In brief, cells were cultured at 37ºC, 5% CO2 in Dulbecco’s Modified Eagle’s Medium (DMEM, Invitrogen Life Technologies, VIC, Australia), supplemented with 10% fetal bovine serum (FBS, Bovigen, VIC, Australia), 1% GlutaMAX and penicillin-streptomcin (Invitrogen). HEK293T cells were then transiently co-transfected with the different voltage-gated calcium channel isoforms and green fluorescent protein (GFP) for visualization, using the calcium phosphate method. cDNAs encoding human Ca_V_3.1 (provided by Dr G. Zamponi), human Ca_V_3.2 (a1Ha-pcDNA3 was a gift from Dr E. Perez-Reyes, Addgene #45809 (Cribbs et al. 1998), human Ca_V_3.3 (a1Ic-HE3-pcDNA3 also from Dr E. Perez-Reyes, Addgene #45810 (Gomora et al. 2002) in combination with green fluorescent protein.

Chinese Hamster Ovary (CHO) cells were used to express K_**V**_1.7 and K_**V**_1.7-Ca_V_3.3 paddle chimeras. Cell culture conditions were the same as the HEK293T cells except DMEM was substituted with DMEM/F12 (Invitrogen). CHO cells were transfected with cDNAs encoding K_**V**_1.7 and K_**V**_1.7/Ca_V_3.3^DI-DIV^ chimeras (synthesized by Genescript, NJ, USA) using Lipofectamine 2000 (Invitrogen) as per manufacturer’s protocol and used for recordings 12-48 hr post transfection.

### Electrophysiology

Whole-cell patch clamp configuration was used to record calcium (I_Ca_) or potassium (I_K_) currents in transiently transfected HEK293T cells. Recordings were made using a MultiClamp 700B amplifier, digitized with a DigiData1440 and controlled using Clampex11.1 software (Molecular Devices, CA, USA). Whole-cell currents were sampled at 100 kHz and then filtered to 10 kHz, with leak and capacitive currents subtracted using a -P/4 protocol for Ca^2+^ currents and uncorrected for K^+^ currents. All recordings were series compensated 60-80%. External solution for I_Ca_ contained in mM: 100 NaCl, 10 CaCl_2_, 1 MgCl_2_, 5 CsCl, 30 TEA-Cl, 10 D-glucose and 10 HEPES, pH 7.3 with TEA-OH. External solution for I_K_ contained in mM: 140 NaCl, 5 KCl, 1 MgCl_2_, 2 CaCl_2_, 10 Glucose and 10 HEPES, pH 7.3 with NaOH. Fire-polished borosilicate (1B150F-4, World Precision Instruments, FL, USA) patch pipettes were used with resistance of 1-3 MΩ. Intracellular recording solution contained as follows (mM): 140 KGluconate, 5 NaCl, 2 MgCl_2_, 5 EGTA and 10 HEPES, pH 7.2 with KOH. Cells were continuously perfused with extracellular solution at a rate of 1.2 ml/min, while toxin application was superfused onto the cell through a capillary tube attached to a syringe pump (2 μl/min), directly onto the cell being recorded. Synthetic μ-Theraphotoxin-Pn3a was kindly provided by the Vetter laboratory (Institute of Molecular Bioscience, University of Queensland) and reconstituted in ddH_2_O.

For experiments on Ca_V_3s, all cells were held at -90 mV. To examine onset of block, test pulses (100 ms, 0.5 Hz) to -20 mV were applied. To generate activation curves, cells were pulsed from - 90 to +40 mV in 10 mV increments at 0.5 Hz. SSI curves were generated by measuring the peak current from a test pulse to -20 mV when preceded by a 1 s pre-pulse from -90 to 20 mV (0.1 Hz). Recovery from inactivation curves were produced by varying the time (0-2 sec) between two depolarizing pules to -20 mV (P1 200 ms, P2 50 ms).

For experiments on K_**V**_1.7 and Ca_V_3.3 DI-IV chimeras, cells were held at -80 mV and activation curves were generated by applying a 50 ms pre-pulse to varying potentials depending of the channel construct examined (K_**V**_1.7 -60 to 90 mV, DI-III 0 to +160 mV, and DIV -120 to +30 mV) followed by a 50 ms test pulse to measure tail currents (K_**V**_1.7 0 mV, DI-III +80 mV, and DIV -20 mV). To measure the onset of Pn3a inhibition, test pulses (50 ms) from a holding potential of -100 mV to a test potential determined for each channel construct (K_**V**_1.7 40 mV, DI-III 120 mV, and DIV 0 mV) were elicited at 0.2 Hz.

### Homology modelling of Human T-type calcium channels and the docking of Pn3a

The comparative model of human T-type calcium channels, Ca_V_3.1, Ca_V_3.2 and Ca_V_3.3 were built upon the Cryo-EM structure of hCa_V_3.1 in the apo form (PDB ID: 6KZO) via SWISS-MODEL server ([*swissmodel*.expasy.org](http://swissmodel.expasy.org)). The 3-D structure of μ-theraphotoxin (TRTX)-Pn3a (PDB ID: 5T4R) was docked into the *three* T-type calcium channels, respectively, via Autodock Vina with a grid box of 40Å × 40Å × 40Å. We also compared the binding mode and the binding affinity among the four VSs of Ca_V_3.3, to identify which VS module Pn3a preferentially targets. The docking results were further analysed via Discovery studio 2017R2 and visualized in visual molecular dynamics (VMD) (Humphrey, Dalke, and Schulten 1996).

### Data and statistical analysis

All data analysis and graphs were generated in OriginPro (Origin Lab Corporation, MA, USA). Concentration-response curves were generated by plotting peak current amplitudes in the presence of Pn3a (I_Pn3a_), over the current prior to Pn3a application (I_Control_). The resulting curve was fit with a sigmoidal curve according to the following expression:

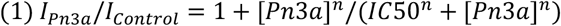

Where IC50 is the half-maximal inhibitory concentration and n is the Hill coefficient. Activation (2) and SSI (3) curves were fit by the modified Boltzmann equation:

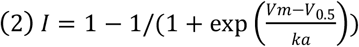

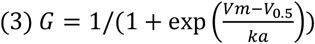

Where I is the current or G is the conductance, Vm is the pre-pulse potential, V_0.5_ is the half-maximal activation potential and ka is the slope factor. Recovery from inactivation plots were fit using a single exponential of the following equation:

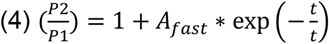

where *τ* is the time constant and *A* is the amplitude. Statistical significance (p < 0.05) was determined using paired or unpaired t-test or 2-way ANOVA followed by a Tukey multiple comparison test if F achieves the level of statistical significance of P < 0.05 and no variance inhomogeneity. All data is presented as mean ± SEM (n), where n is individual cells with all experimental results containing n ≥.5 individual cells.

## Acknowledgments

This work was supported by the Rebecca Cooper Foundation for Medical Research Project Grant [PG2019396] to J.R.M and the National Health and Medical Research Council (NHMRC) Program Grant [APP1072113] to D.J.A.. J.R.M and R.K.F-U are grateful to Emma and Zack O. Yepugas for continuous support. Computational resources were provided by the National Computational Infrastructure (NCI) which is funded by the Australian Government, and the Pawsey Supercomputing Centre which is funded by the Australian Government and the Government of Western Australia.

---

**Figure 3-figure supplement 1.**
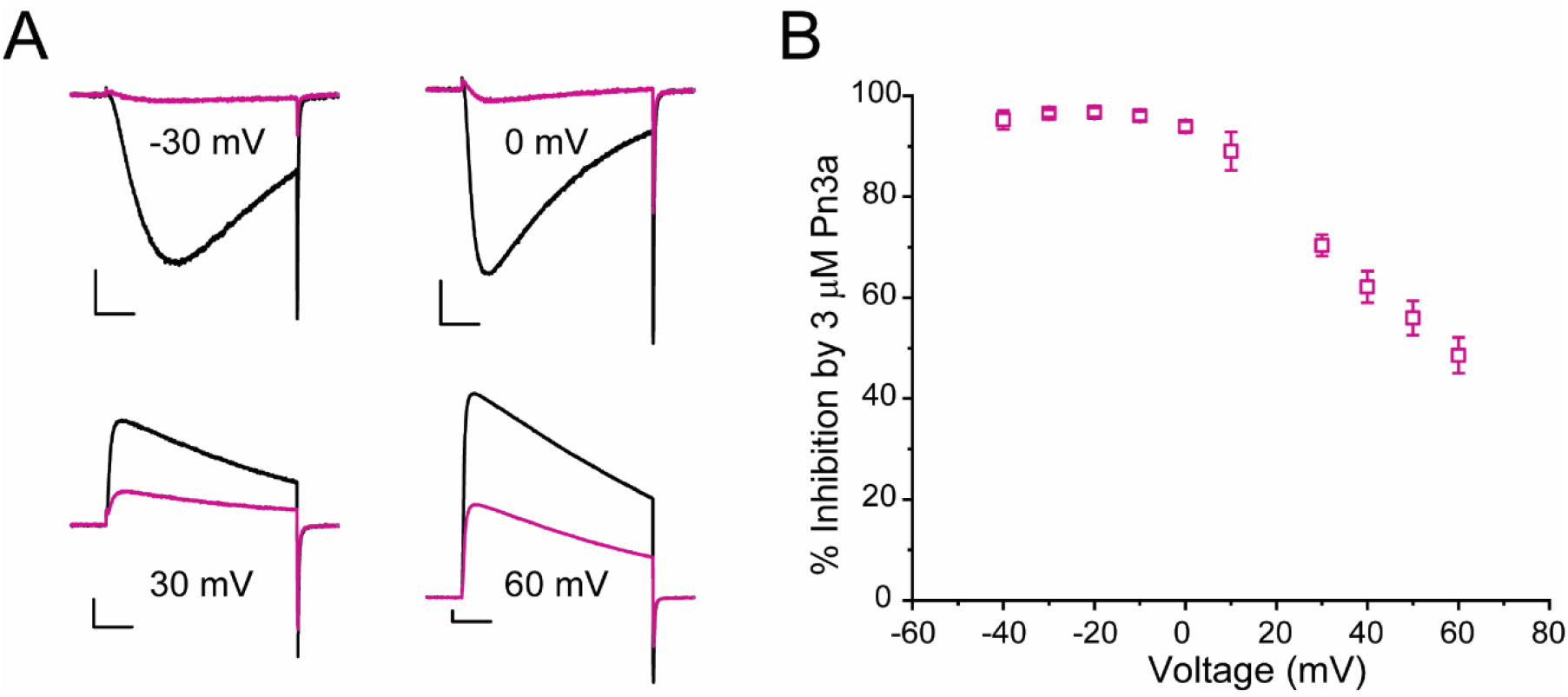
Voltage dependence of Pn3a inhibition of human Ca_V_3.3 when Na^+^ is the charge carrier. **(A)** Representative traces showing the inhibition by 3 μM Pn3a of inward and outward I_Na_ through Ca_V_3.3 at −30, 0, 30 and 60 mV. Scale bars: 1 nA, 20 ms. **(B)** Graph showing % inhibition of I_Na_ by 3 μM PnAa from −40 to +60 mV.

**Figure 6-figure supplement 1.**
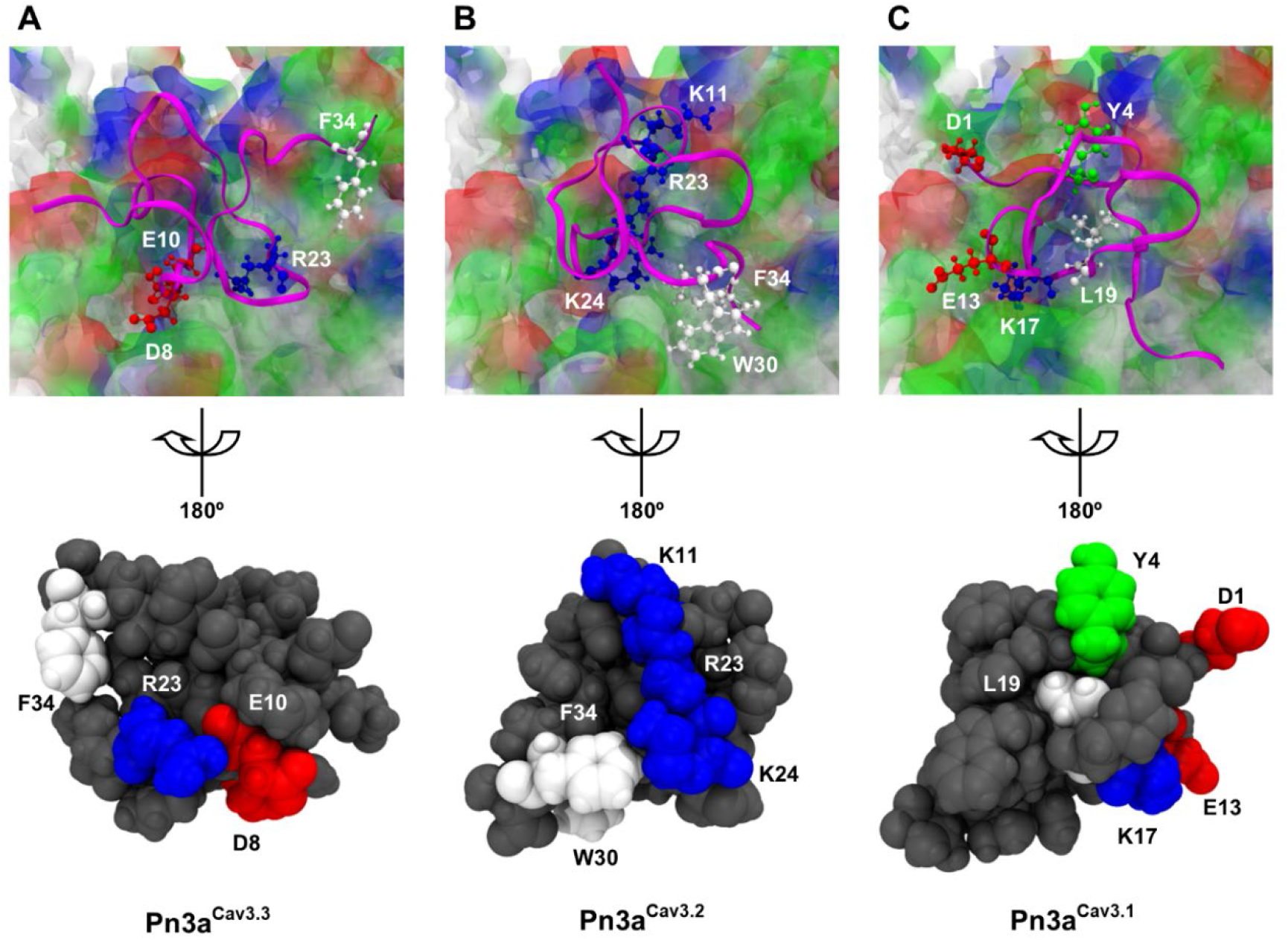
The binding conformation of Pn3a at voltage-sensing domain (VSD)-II of T-type calcium channels. The interactive residues of the spider toxin contacting with corresponding residues of the VSDII of Ca_V_3.3 (A), Ca_V_3.2 (B) and Ca_V_3.1 (C), respectively. For clarification, the binding domain of Ca_V_3 channel shown in transparent color was removed after rotating the binding conformation around the y-axis by 180º (in the bottom). The coloring method in VMD was adopted for the residues and surfaces, demonstrating non-polar (white), basic (blue), acidic (red) and polar (green) residues.

**Figure 6-figure supplement 2.**
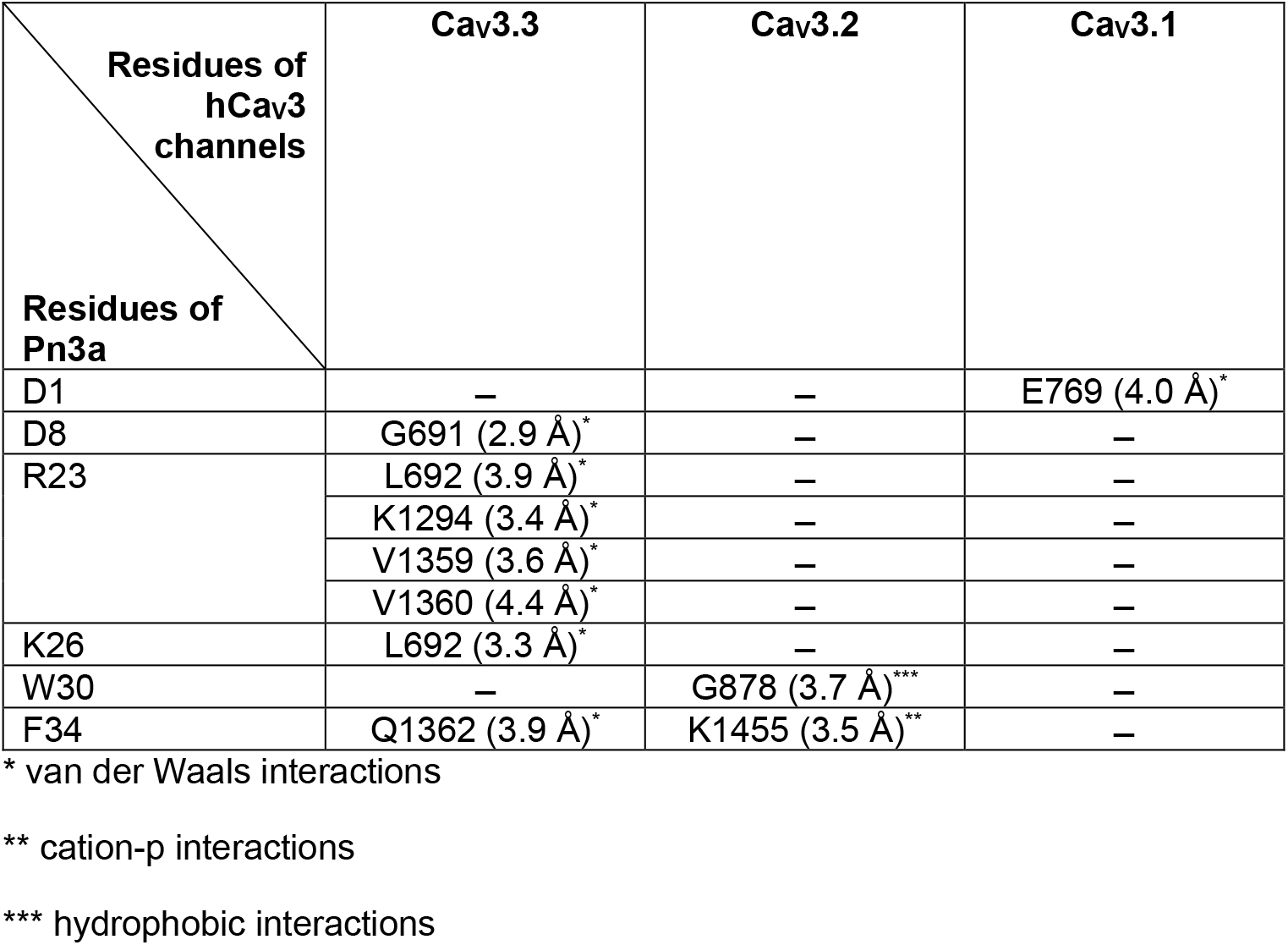
Minor interactions between the residues of Pn3a and the corresponding residues on hCa_V_3 channels.

